# Machine-driven parameter optimisation of biochemical reactions

**DOI:** 10.1101/739771

**Authors:** Stéphane Poulain, Ophélie Arnaud, Sachi Kato, Iris Chen, Hiro Ishida, Piero Carninci, Charles Plessy

**Affiliations:** RIKEN Center for Life Science Technologies, Division of Genomics Technologies, Yokohama, Japan; RIKEN Center for Integrative Medical Sciences, Division of Genomic Medicine, Yokohama, Japan; Biomedical Microsystems Lab., Institute of Industrial Science, The University of Tokyo, Japan; Université Claude Bernard Lyon 1, INSERM 1052, CNRS 5286, Cancer Research Center of Lyon, Centre Léon Bérard, F-69008, Lyon, France; Labcyte Inc., Tokyo, Japan; Okinawa Institute of Science and Technology Graduate University, Genomics and Regulatory Systems Unit, Onna-son, Japan

## Abstract

The development of complex, multi-step methods in molecular biology is a laborious, costly, iterative and often intuition-bound process where an optimum is sought in a multidimensional parameter space through step-by-step optimisations. The difficulty of miniaturising reactions under the microliter volumes usually handled in multiwell plates by robots, plus the cost of the experiments, limit the number of parameters and the dynamic ranges that can be explored. Nevertheless, because of non-linearities of the response of biochemical systems to their reagent concentrations, broad dynamic ranges are necessary. Here we use a high-performance nanoliter handling platform and computer generation of liquid transfer programs to explore in quadruplicates more than 600 combinations of 4 parameters of a biochemical reaction, the reverse-transcription, which lead us to uncover non-linear responses, parameter interactions and novel mechanistic insights. With the increased availability of computer-driven laboratory platforms for biotechnology, our results demonstrate the feasibility and advantage of methods development based on reproducible, computer-aided exhaustive characterisation of biochemical systems.

## Introduction

Systematic explorations of reaction parameters have been driven by automation and miniaturisation of laboratory experiments, and sub-microliter liquid handling systems hold the biggest promises in terms of throughput together with reducing the cost of reagents. For instance, microfluidics technologies were used by Kim *et al.* (2011) to generate 64 different combination of salt and DNA concentrations in a hybridisation assay, for 5 different salts. More recently, Genot *et al*. (2016) measured thousands of unique combinations of different concentrations of reagents and their reaction products using microfluidic droplets. However, such approaches rely on merging physical streams of reagents, and thus are limited in the number of parameters that can be explored simultaneously and are also hard to apply to categorical parameters. On the other hand, several nanoliter-handling platforms have appeared on the market and empower researchers to design digitalised analysis with an arbitrary number of reagents. Among these platforms we chose acoustic droplet ejection technology because it combines several advantages. First, there is no contact between the machine and the liquid, which eliminates the cost of disposable plastic pipette tips. Second, source reagents can be provided in microplates with hundreds or even thousands of wells, allowing for the use of molecular barcodes. Third, each liquid transfer is fast.

As a pilot reaction for optimising, we chose reverse-transcription, which converts messenger RNA (mRNA) molecules to complementary DNA (cDNA), a suitable substrate for quantitative DNA sequencing technologies (Bentley *et al*., 2008). The reverse-transcription reaction is central to biological experiments that aim at quantifying the activity of genes. In experiments where the amount of biological substrate is limited in quantity, the performance of the reverse-transcription reaction becomes a limiting factor. In particular, analysis of single cells requires a highly efficient conversion from mRNA to cDNA (Zucha *et al.*, 2019). The reverse-transcriptase enzyme needs a short DNA oligonucleotide to prime the reaction. In addition, many methods for single-cell analysis use an additional “template-switching” oligonucleotide in order to extend the cDNA sequence in preparation for sequencing (see Figure 1 and for review, Picelli, 2017). While increasing concentrations of the reverse-transcription primer and the template-switching oligonucleotide tend to increase the efficiency of the reaction, a limit is reached at high values (see for instance Zajac *et al.*, 2013). The proliferation of protocols using clearly different reaction parameters (Table 1) suggests rather that optimal oligonucleotide concentrations vary with other parameters such as enzyme type or substrate amounts.

**Table 1.** Comparison of reverse-transcription reagents concentrations used in different RNA sequencing protocols. TSO: template-switching oligonucleotide. RTP: reverse-transcription primer.

| TSO (nM) | RTP (nM) | dNTP (mM) | Mg (mM) | Mn | DTT | Enzyme (units) | Reference |
| --- | --- | --- | --- | --- | --- | --- | --- |
| 10,000 | 1000 | 2.5 | 3.0 | 0 | 10 | SSIII (200) | Poulain 2017 |
| 1,000 | 1000 | 2.0 | 7.2 | 0 | 4 | SSII (50) | Lee 2017 |
| 4,600 | 135 | 1.2 | 8.7 | 0 | 5.4 | SSII (6.8) | Hochgerner 2017 |
| 1,000 | 100 or 1 | 1.0 | ? | 0 | 2.4 | SmartScribe (100) | Turchinovich 2014 |
| 1,000 | 1000 | 1.0 | 12.0 | 0 | 2.5 | SSII (100) | Picelli 2013 |
| 200 | 200 | 1.0 | 3.0 | 3 | 1 | SSII (5) | Islam 2011 |
| 0 | 750 | 4.0 | 3.0 | 2 | 5 | SSII (200) | Schmidt 1999 |
| 0 | 65789 | 1.1 | 3.0 | 0 | 5 | SSIII (760) | Murata 2014 |

**Figure 1:**
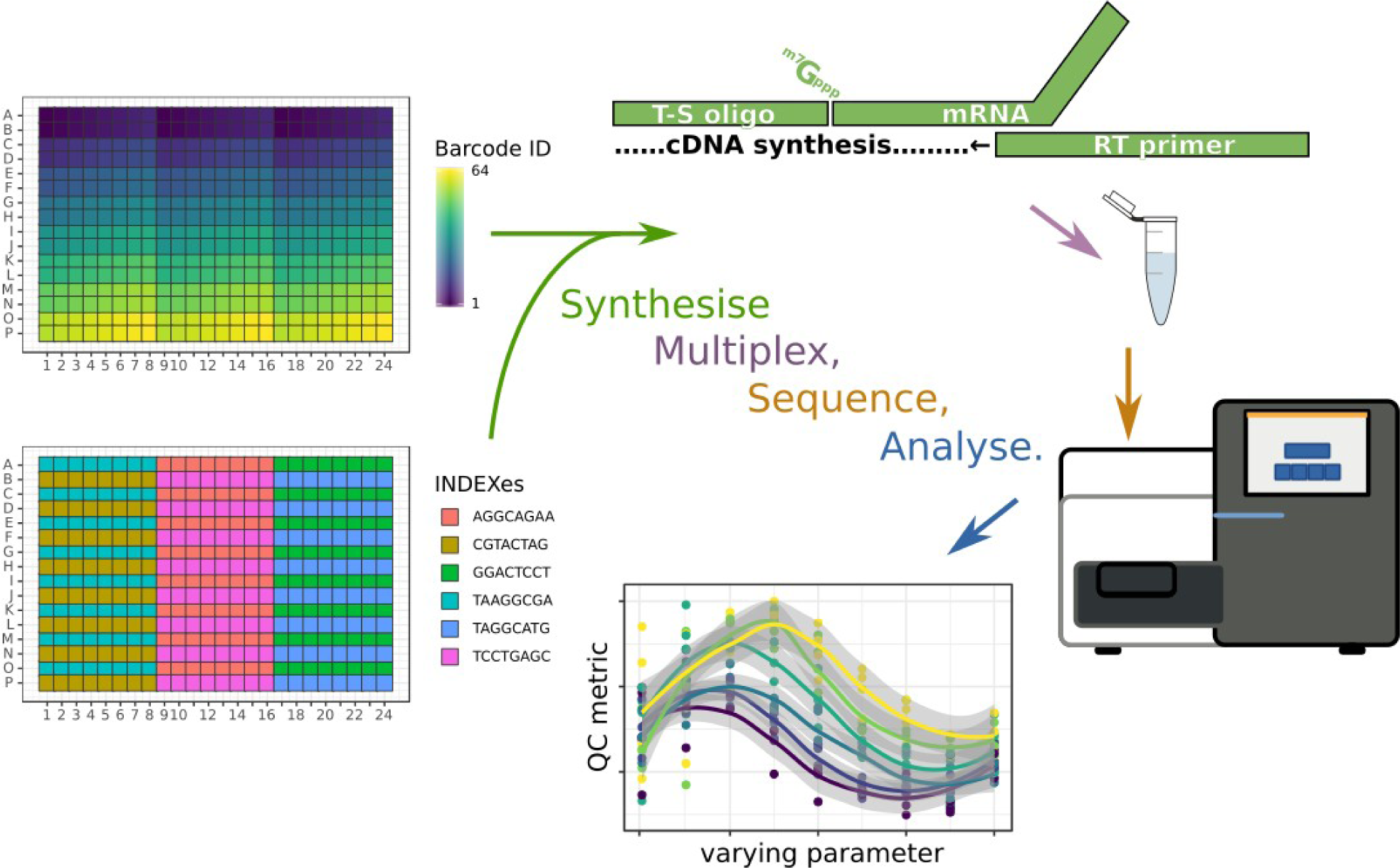
Method overview. This conceptual drawing shows a 384-well plate where each individual well is recognised by a combination of 64 different “barcodes” and 6 different “indexes”. Wells with the same index are interleaved to match the geometry of standard multichannel pipettes and collected together in the same microtube. The studied reaction is a reverse-transcription where a mRNA is converted to cDNA by a reverse transcriptase (RT) using a primer. In addition, a template-switching (T-S) oligonucleotide is present in the reaction to extend the sequence of the mRNA. After incubation in the plates, reactions prepared with the same starting amounts of RNA are amplified together in six different polymerase chain reactions using indexed primers, pooled in a multiplexed sequencing reaction, and then demultiplexed in silico using the unique combination of barcode and index sequences. Multi-factorial quantitative analysis of the sequencing reads then follows, according to the initial reaction parameters decided for each well in the experiment design.

In the past decade, we have developed a method for quantitative high-throughput gene expression analysis, nanoCAGE (Plessy *et al.*, 2010; Poulain *et al.*, 2017), in which a key step is a template-switching reverse-transcription. The desired product of a nanoCAGE reaction is a set of sequence reads that align in gene promoters at transcription start sites in proportional quantities to a gene’s expression level. Several indicators are used to assess the quality of a nanoCAGE reaction. “Oligonucleotide artefacts” are sequences that match synthetic DNA adapter sequences that should not be found within the reads, such as PCR primer dimers. The “ribosomal rate” is the proportion of the reads aligning to ribosomal RNA sequences. Since ribosomal RNAs are very abundant, their sequencing is at the expense of the sequencing of other genes and therefore must be avoided to reduce the experimental cost. After removal of the ribosomal RNA sequences and other low-complexity sequences, the “mapping rate” is the proportion of reads that could be aligned to a reference genome and measures the amount of sequencing reads that is effectively spent measuring gene expression levels. The “strand invasion rate” is the number of reads suspected to be produced by a specific artefact of the template-switching reaction. These artefacts, detected after mapping, cause mischaracterisation of promoters and bias the measurement of their activity (see Tang *et al.*, 2013 for details). Lastly, after removal of the strand-invasion artefacts, the “promoter rate” is the proportion of reads aligning to promoter regions and therefore likely to indicate true transcription start sites. These quality control metrics converge quickly, requiring less than a thousand of reads per experiment, therefore allowing us to massively multiplex nanoCAGE reactions in small-scale, cost-efficient sequencing runs. Using them as a proxy for determining the quality of reverse-transcription reactions, we searched for an optimum by a systematic exploration of the parameter space.

## Results

Using a Labcyte Echo 525 instrument, we assembled 500 nL reverse-transcription reactions in 384-well plates, by dispensing droplets of 25 nL from a source plate containing the reagent stocks to the destination plates. After incubating the reactions, we assessed their yield and quality by quantitative DNA sequencing using the nanoCAGE method. We multiplexed two pools of reactions (2 × 1,536) using a selected set of 64 different template-switching oligonucleotides carrying sample identifiers (“barcodes”), combined with 24 sequencing indexes (Figure 1). As we focused on quality controls, the sequencing was kept at a small scale: ∼5,000 sequence read pairs per reaction.

We explored one categorical and three continuous parameters spanning multiple orders of magnitude with data points regularly spaced on a logarithmic scale. The broadest range was for RNA amounts, with 6 points ranging between single-cell amounts (1∼10 pg) and starting amounts typical for bulk RNA libraries (10∼100 ng, or a few µg in larger reaction volumes). The second broadest range was the molarity of the template-switching oligonucleotide (9 points between 0.6 and 160 µM), followed by the reverse-transcription primer’s molarity (6 points between 1 and 24 µM). Combined with a categorical dimension encoding 2 different reverse-transcription enzymes: SuperScript III (SSIII) and SuperScript IV (SSIV), we thus generated 648 different combinations, which we studied in quadruplicates. We added 120 negative control reactions per replicate, for a total of one 384-well plate per enzyme and replicate.

To prevent position biases in the reaction microwell plates, such as drying on the edges or inhomogeneous heating during reaction, we randomised the coordinates of the reactions in each replicate. Therefore, a given combination of parameters will not appear twice in the same plate position, except by chance. This randomisation also has the effect of making each replicate plate unique, which prevents from accidental swapping. In particular, the different positions of the negative controls in each plate act as an identifying fingerprint. Lastly, since we used barcoded oligonucleotides, the randomisation prevented that barcode sequences, individual reagent quality, or position-specific variability in the reagent transfer would get confounded with any reaction parameter. Randomisations were made entirely reproducible by the computer-aided generation of machine-readable volume transfer instructions and were reproducibly repeated for each replicate with a different random seed.

To ensure equal sequence coverage, the PCR amplifications that followed the reverse-transcriptions were scaled according to the starting quantities of RNA. Each PCR amplification was multiplexed with a different index (Figure 1A). For each sample amplified in the same PCR reaction, we then calculated relative yields, and observed a positive correlation with the molarity of the template-switching oligonucleotide (Figure 2A), and little effect of the reverse-transcription primer, except at the highest concentrations of both oligonucleotides and the RNA (Figure 2B). To better represent the effect of the interaction between the two oligonucleotides, we calculated the medians of each replicates, and represented them as a contour plot (Figure 2C) on surfaces where the molarity of each oligonucleotide is one dimension, and with an arbitrary colour scheme where blue indicates preferable results (here, higher yields). We use the same graphical representation for other quality control statistics in the remaining figures of the manuscript. As these plots do not show directly the variability between replicates, categorical plots versions in the same style of Figure 2B are available in the supplementary material.

**Figure 2:**
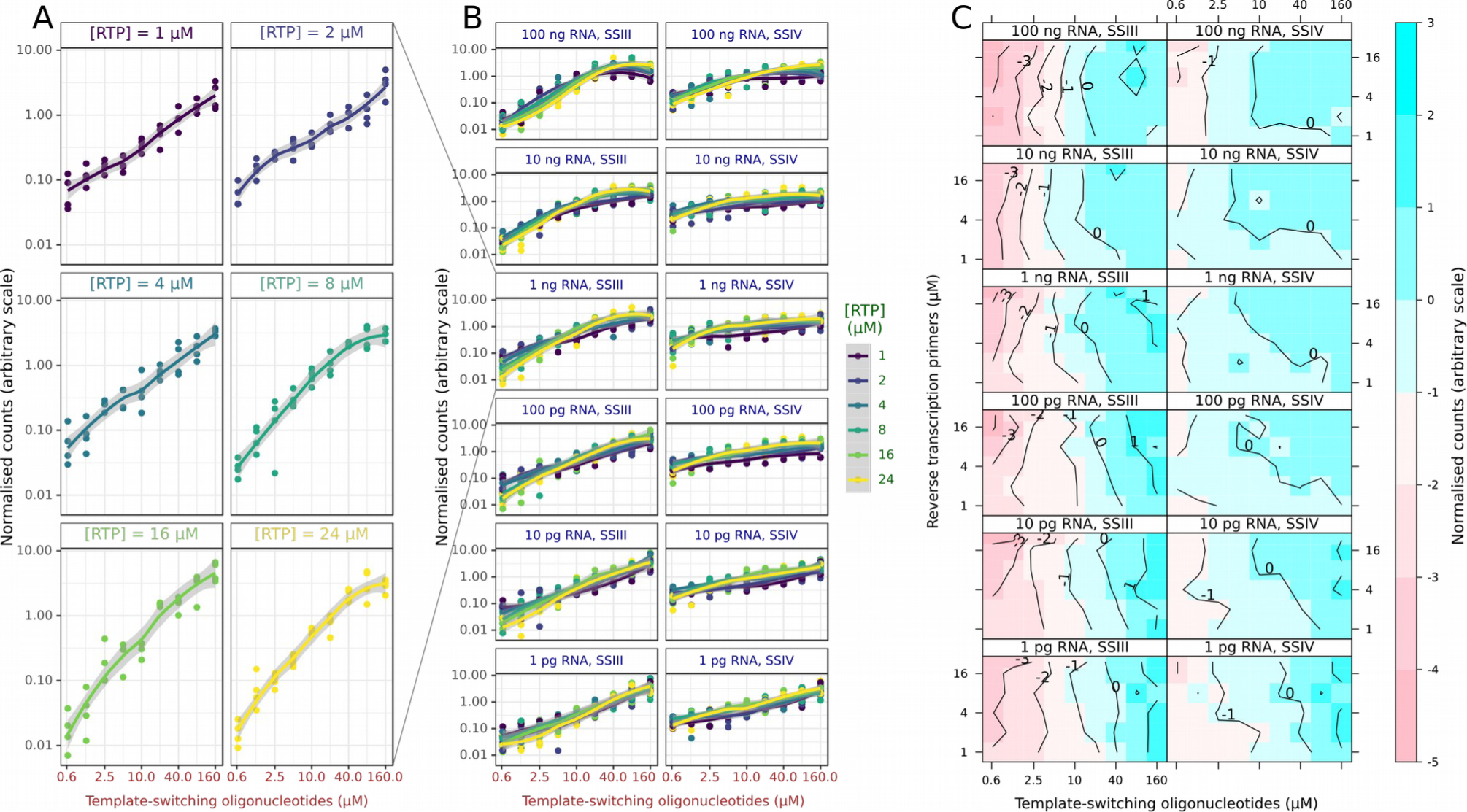
Parameter screen at high dynamic range. A: log-normalised reaction yields as a function of the molarity of the template-switching oligonucleotide, for different molarities of the reverse-transcription primers (RTP), using the SSIII enzyme and 100 pg RNA. B: same data displayed for all RTP molarities, colour-encoded as a categorical variable, and the SSIII and SSIV enzymes. In this panel, each replicate is displayed as a dot. C: Median yield over each set of four replicates, displayed as a contour plot on a surface where each axis represents the molarity of one oligonucleotide.

To assess the quality and specificity of the reactions, we then calculated the quality metrics that were described in the introduction (Figure 3). With RNA inputs lower than 100 pg, the SSIII enzyme started to produce large amounts of oligonucleotide artefacts, at a scale that would compromise its use in transcriptome analyses. The SSIV enzyme performed comparatively well on one order of magnitude lower RNA input (10 pg). For both enzymes, increasing the molarity of the reverse-transcription primer lead to an increase of the oligonucleotide artefacts, but this could be compensated by a proportional increase of the molarity of the template-switching oligonucleotide, as shown by the diagonal patterns on the contour plots (Figure 3A). A similar diagonal pattern can be observed in the contour plots for the ribosomal RNA rate (Figure 3B), showing again a positive effect in terms of quality for the reduction of the reverse-transcription primer’s molarity and the increase of the template-switching oligonucleotide, especially for the SSIV enzyme. Another interesting trend was that overall, ribosomal RNA rates were lower for two extreme amounts of starting RNA material (100 ng and 1 pg). For SSIII at all RNA amounts and for SSIV at low RNA amounts, the highest molarities of the template-switching oligonucleotide were also increasing the ribosomal RNA rate. We then calculated the mapping rate, in which the ribosomal RNA sequences are considered unmapped. The contour plots (Figure 3C) were essentially the reverse of Figure 3B, showing that most non-ribosomal reads mapped correctly.

**Figure 3:**
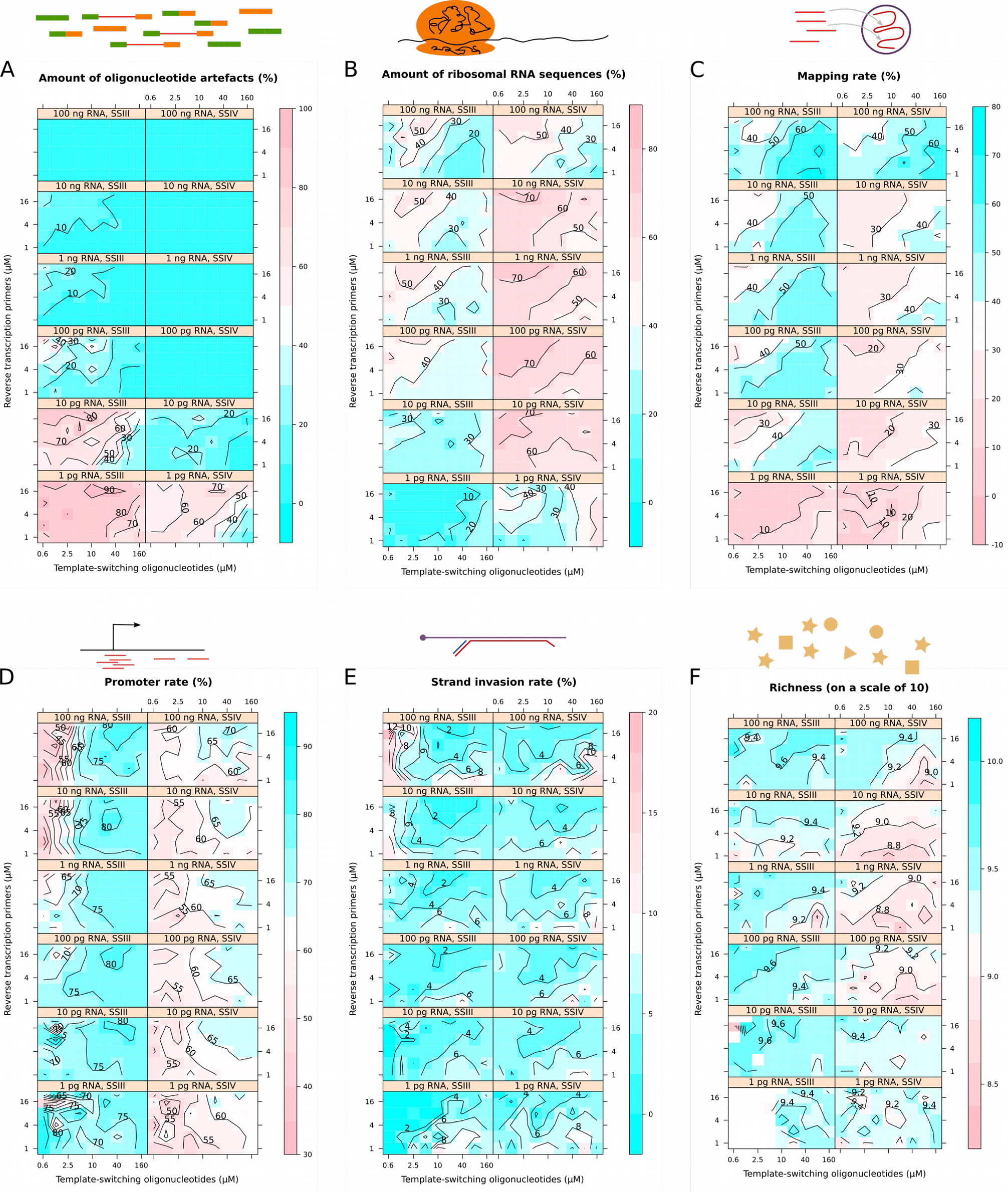
quality and specificity across the parameter space. proportions of (A) oligonucleotide artefacts, (B) reads mapping to reference ribosomal RNA sequences, (C) reads mapping to the genome, (D) reads aligning to promoter regions, (E) premature “strand invasion” artefacts of the template switching reaction and (F) richness index.

Within the set of all mapped reads, we calculated the promoter rate (Figure 3D). Both enzymes showed similar trends except that SSIV had a lower baseline. Increases of the molarity of the template-switching oligonucleotide again increased reaction quality except at the highest molarities. Strikingly, the best promoter rates were obtained at high molarity of the reverse-transcription primer, which is the opposite trend in comparison with the other quality metrics calculated above. All RNA amounts benefited from high oligonucleotide molarities.

We then calculated the proportion of mapped reads that could be strand-invasion artefacts (Figure 3E), which form when the template-switching oligonucleotide prematurely hybridises with the nascent cDNA (Tang *et al.*, 2013). Contrarily to the intuition that higher concentrations of this oligonucleotide would lead to higher frequencies of artefacts at higher molarity of the template-switching oligonucleotide, our results show a more complex relationship. For instance, the most extreme strand-invasion rates were reached with SSIII at high RNA amounts and low template-switching oligonucleotide molarity. This suggests that under these conditions, the artefacts may be more easily created than the desired products, and that higher concentrations of template-switching oligonucleotide and reverse-transcription primer are necessary to repress artefact formation. Thus, our approach gave us not only a fine-grained mapping of the optimal reaction conditions in the parameter space, but also mechanistic insights.

While our strategy of shallow sequencing allows us to calculate accurate quality metrics despite a very low coverage of each reaction, it is impossible to compare the expression profiles with each other, or to know the total number of genes that would be detected if we had orders of magnitude deeper coverage. To estimate the potential for gene detection, we calculated richness indexes for each library following the method of Hurlbert (1971), on a scale of 10 (Figure 3F). The resulting numbers answer the question “how many genes would we expect to detect if we sequenced 10 reads randomly from this library?”. Despite being on an arbitrary scale, richness indexes are very useful to transcriptome analysis as they converge very quickly, even at depth where most of the expressed genes are not yet detected. In our libraries, surprisingly, richness indexes were higher at lower template-switching oligonucleotide molarities and higher reverse-transcription primer molarities. While intuitively it seems desirable to have higher richness indexes, it remains to be determined if they might be the reflection of a compression of the dynamic range.

Our results confirm that bulk and single-cell reactions have different optima. At high amounts of starting RNA (100 ng), the standard nanoCAGE protocol (SSIII enzyme, 10 µM template-switching oligonucleotide, 1 µM reverse-transcription primer) is close to the optimal results, minimising the amount of oligonucleotide and ribosomal RNA artefacts and maximising the amount of data that correctly aligned to promoter regions. On the other hand, our results suggest that a protocol can be designed for single cells by using the SSIV enzyme, increasing the molarity of template-switching oligonucleotides and reducing the molarity of the reverse-transcription primers. Indeed, we validated this observation during the development of a single-cell version of the nanoCAGE protocol, where we tested the use of SSIV with the template-switching oligonucleotide molarity increased to 45 µM and the reverse-transcription primer molarity decreased to 0.4 µM, on single cells isolated by flow cytometry in 4 µL reverse-transcriptions reactions. As the amount of RNA per cell is usually estimated to be in the order of magnitude of 10 pg, the reaction concentration is in the order of the 1 pg / 500 nL condition in our parameter screen. In line with the screen’s results, the change of enzyme and oligonucleotide concentrations dramatically reduced the amount of oligonucleotide artefacts, although at the expense of a reduced promoter rate and an increased rRNA rate (Figure 4).

**Figure 4:**
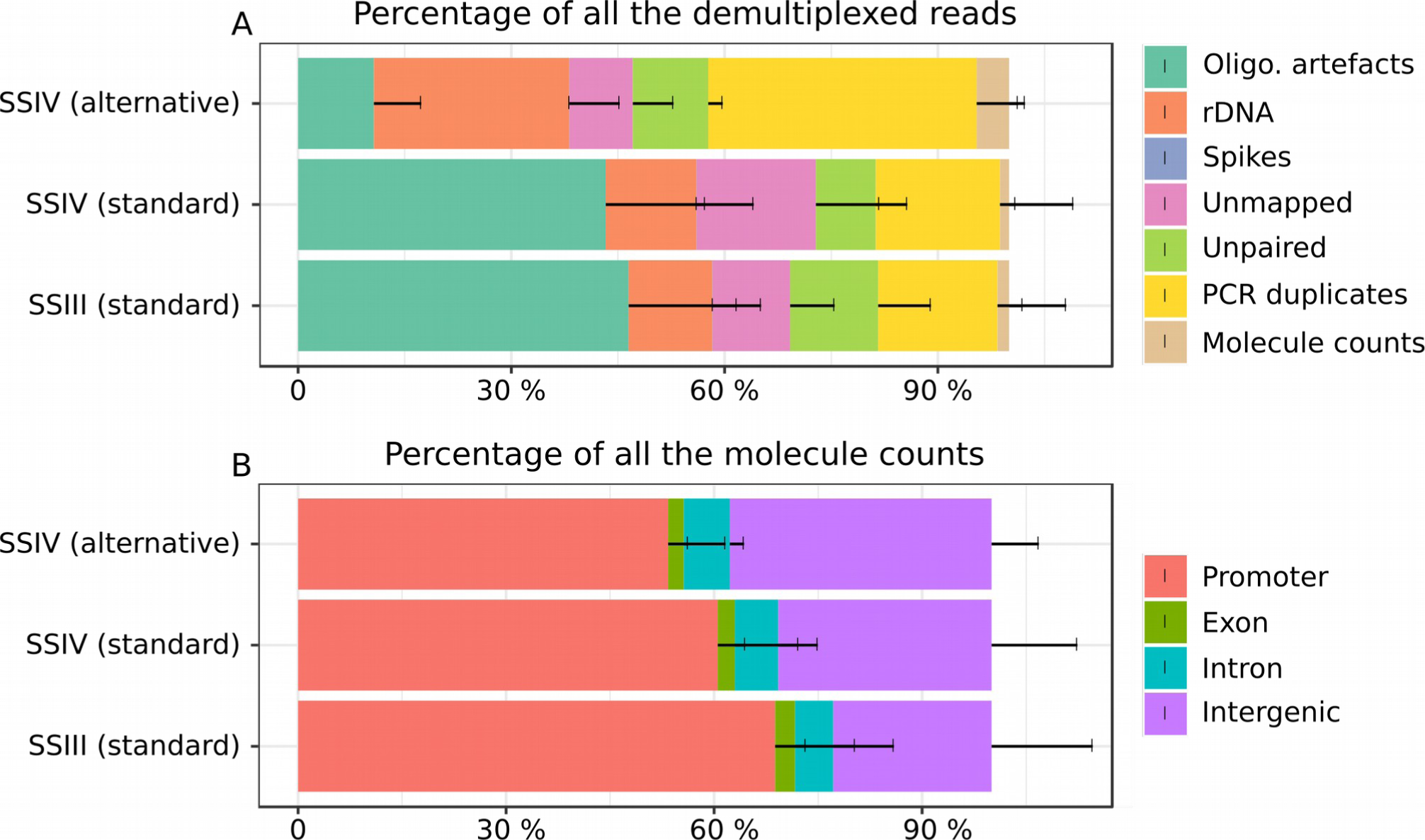
Test of alternative protocol on low RNA inputs (single cells). Stacked barplots summarising the quality control statistics for single cell libraries made with the standard nanoCAGE protocol (SSIII standard, n=136), or with the SSIV enzyme and the standard oligonucleotide concentrations (SSIV standard, n = 184), or with the SSIV enzyme with 45 µM template-switching oligonucleotide and 0.4 µM reverse-transcription primer (SSIV alternative, n=83). A: proportions of the reads discarded during data processing until obtaining unique molecule tag counts. B: annotation statistics of the aligned molecule tags.

## Discussion

To our knowledge, we are the first to report the exploration of a four-dimension parameter space for biochemical reaction using 648 data points. Our experimental design described here follows a systematic and exhaustive approach testing all combinations (“grid search”) between a few parameters. Nevertheless, such a design cannot be extended to a larger number of parameters, because their total number of combinations would not fit into 384-well plates (although it would take advantage of smaller formats such as 1536-well plates or miniaturised array), and because high-dimension spaces are typically sparse. More efficient search strategies, such as random search (Bergstra & Bengio, 2012), may enhance the power of the method.

We observed a dramatic difference between the two reverse-transcriptases. According to the maker, SSIV has better thermostability, processivity and yield that SSIII. However, it is important to note that their reaction buffers also differ: SSIV’s buffer is proprietary and is advertised to perform better in the presence of inhibitors. Swap experiments would be needed for determining whether the enzyme or the buffer are the key factor driving our observations. Another simple explanation could be that the unit definition, which is “the amount of enzyme required to incorporate 1 nmole of deoxyribonucleotide into acid-precipitable material in 10 min at 37°C using poly(A) oligo(dT)_12-18_ as a template/primer” may correspond to different actual quantities of enzyme, given their different thermal optima. Beyond addressing such questions, the approach that we have developed here will also provide the opportunity to search for ad-hoc mixtures of enzymes, and possibly buffers, that may maximise the output of the reaction (conversion to cDNA) and its quality (efficiency and specificity of template switching).

The reverse-transcription reaction that we studied here uses so-called “random” primers that end with six random bases, for an even coverage of the transcriptome. Their drawback is the creation of artefacts when they hybridise to other oligonucleotides. In line with this, we observed that increasing the molarity of these reverse-transcription primers was increasing the amount of artefacts in the sequencing libraries (Figure 3A). Unfortunately, an opposite trend was seen on other quality statistics, such as the promoter rate (Figure 3D). This conflict may be hard to resolve. There are alternatives to random primers, such as oligo-dT primers that target the poly-A tail of the mRNAs, not-so-random primers (Armour *et al.*, 2009) or pseudo-random primers (Arnaud *et al.* 2016). While the discussion of their pros and cons is beyond the scope of this manuscript, it is important to note that their optimal molarities are likely to be different.

Methods in molecular biology are made of a large number of serial steps, and the approach that we followed here is not limited to the reverse-transcription reaction. Here, we used quantitative sequencing for the readout, and this strategy can also apply to other reactions. For instance, the activity of a DNA-cutting enzyme can be assayed by the degradation of a sequencing template. Conversely, the activity of a DNA-joining enzyme can be assayed by the assemblage of a template. Other platforms for the readout can be used, such as fluorescence detection, obviously for quantitative polymerase chain reaction, but also for any other reaction that can be designed to produce or degrade a fluorescent reporter.

Beyond the example presented here as a proof of principle, we believe that parameter space screening may become a routine experiment in the future. In particular, it will greatly benefit from computer-aided experiment design and robotic automation of experiment execution (Yachie *et al*., 2017, McClymont *et al*., 2017). The combination of laboratory automation and systematic parameter screening will ease the way to cross-replication studies in independent laboratories, as a strategy for cost-sharing, removal of implementation bias, and detection of human errors or data tampering.

## Methods

Nanoliter-scale liquid transfers were performed on an Echo 525 instrument (Labcyte). The transfer sheets were generated from event logs produced by simulations of instrument runs in the R programming language using the layout of source and destination plates as input. All the scripts are available on https://gitlab.com/charles-plessy/labcyte-rt-optimisation.

We used a total RNA prepared from mouse liver. An RNA Integrity Number (RIN, Schroeder *et al.*, 2006) of 9 was measured on an Agilent Bioanalyzer with an RNA 6000 pico kit. As molarity measurement were not possible with this kit, RNA amounts are measured as a mass in this work.

The template-switching oligonucleotides were synthesized by Integrated DNA Technologies, Inc. (IDT), at a scale of 10 nmoles each in a 96-well plate format and resuspended at a stock concentration of 1 mM upon reception. To assess the quality of synthesis, we ran a control experiment in which the parameters of the reverse-transcription reactions were kept constant (see supplementalexperimentsixon https://gitlab.com/charles-plessy/labcyte-rt-optimisation/blob/master/Labcyte-RT_Data_Analysis_6.md for details). For the main experiment, we selected 64 oligonucleotides among those who gave yields closest to the mean.

RT reaction master mixes (containing 0.0528 M sorbitol, 0.264 M trehalose, 0.75 M betain, 0.01 M DTT, 0.625 mM dNTPs, 20 U/µL SuperScript enzyme III or IV and 1× SuperScript buffer III or IV), template-switching oligonucleotides, reverse-transcription primers, RNA and ultrapure water were transferred from independent wells of a source plate (Greiner Bio-One) to the different wells of the destination plates (Applied Biosystems) using specific liquid transfer patterns for each plate. After the acoustic transfer of RT reagents (total RT reaction volume equal to 500 nL), the 384-well destination plates were directly sealed, centrifuged at 4 °C, and deposited in a 7900HT Fast Real-Time PCR system (Applied Biosystems) to perform first strand cDNA synthesis (temperature program: 22 °C for 10 min; 50 °C for 30 min; and 70°C for 15 min). RT products were further collected with a multichannel pipette, pooled and purified using AMPure XP beads (Beckman Coulter). Purified pools were then processed as in the nanoCAGE protocol (Poulain et al., 2017) applying 18, 21, 24, 27, 30 and 33 PCR cycles during the first amplification reaction, respectively for pools of RT reactions corresponding to template RNA amounts ranging from 100 ng to 1 pg. Libraries prepared for each starting RNA amount were tagged by specific index sequences (Nextera XT DNA Library Preparation Kit, Illumina), mixed equimolarly and sequenced on a MiSeq system using a reagent kit v2 (Illumina). Sequencing data were processed with the MOIRAI workflow system (Hasegawa *et al.*, 2014) (Supplementary Figure 3) to produce alignment files (BED) and sequencing quality metrics that were analysed using custom R scripts.

**Supplementary Figure 3:**
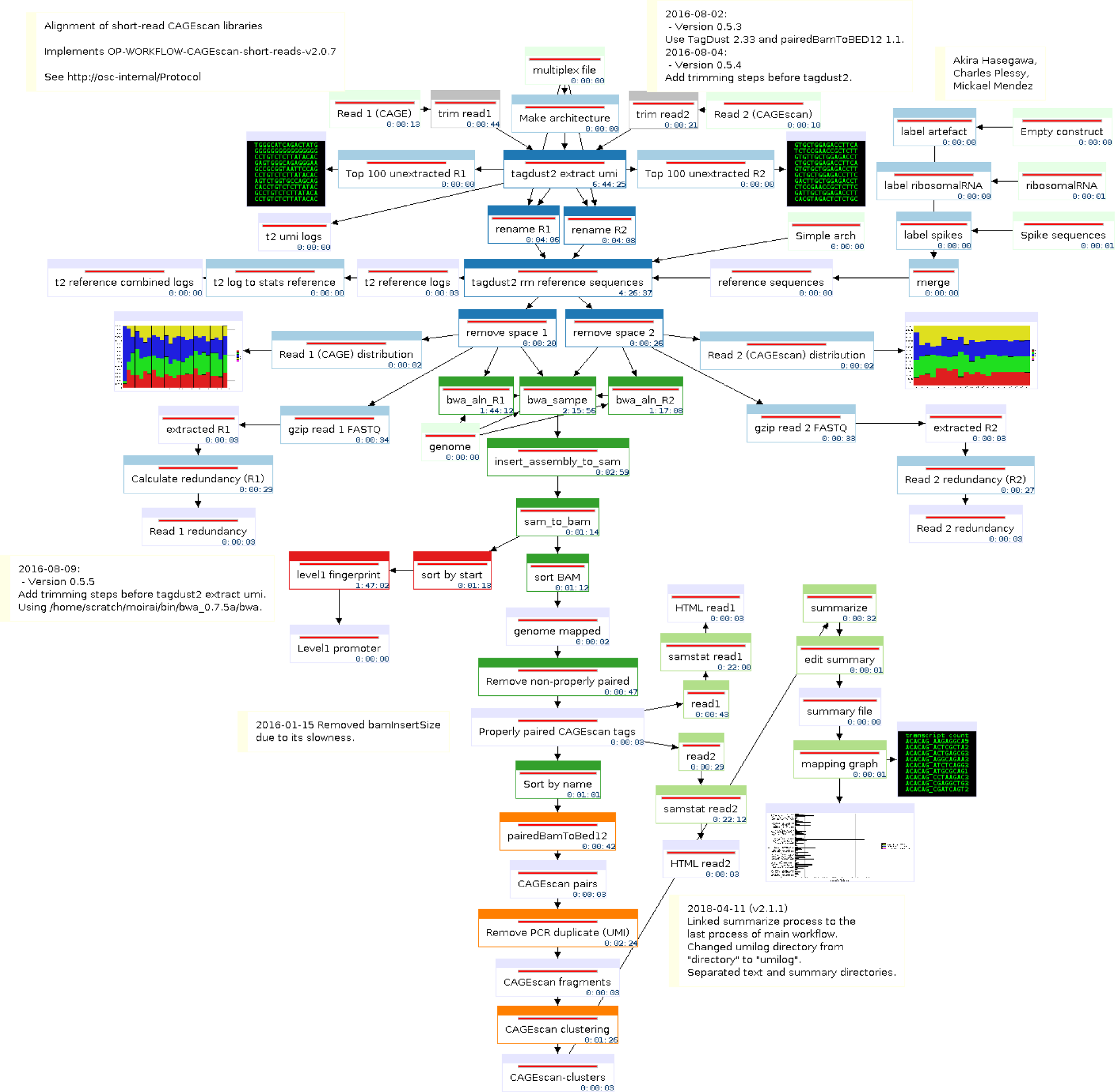
graphical summary of the MOIRAI workflow used to process the sequencing data. The full data for each sequencing run has been deposited to ZENODO (https://doi.org/10.5281/zenodo.1683162).

For single cell experiments, a suspension of HeLa cells (ATCC ref. CCL-2) was prepared, filtered through a 40 µm cell strainer (Falcon), and sorted by flow cytometry (FACSAria II, BD Biosciences) in order to transfer individual cells into 96-well plates (Applied Biosystems). After single cell isolation, the plates were directly sealed, centrifuged at 4 °C, and stored at –80 °C until further processing. Cell lysis was initiated upon thawing frozen plates on ice for 15 min. Next, RT master mixes (containing 0.0528 M sorbitol, 0.264 M trehalose, 0.75 M betain, 0.01 M DTT, 0.625 mM dNTPs, 1 µM or 0.4 µM RT primers, 20 U/µL SuperScript enzyme III or IV and 1× SuperScript buffer III or IV) were transferred with a multichannel pipette. Specific barcoded template-switching oligos (10 µM or 45 µM) were then added in each well according to the multiplexing plan. After the addition of RT reagents (total RT reaction volume equal to 4 µL), the plates were directly sealed, centrifuged at 4 °C and placed in a 7900HT qPCR system to carry out first strand cDNA synthesis (temperature program for SuperScript IV: 22°C for 10 min; 50°C for 15 min; and 80°C for 10 min; temperature programm for SuperScript III: 22°C for 10 min; 50°C for 30 min; and 70°C for 15 min). RT products were harvested with a multichannel pipette, pooled and purified using Ampure XP beads (Beckman Coulter). Purified pools of single cell RT products were processed for nanoCAGE library preparation (Poulain et al., 2017). Pools of reactions performed using standard primer concentrations with SusperScript III or SuperScript IV; and using alternate primer concentrations with SuperScript IV were subsequently tagged by specific index sequences (Nextera XT DNA Library Preparation Kit, Illumina), pooled and sequenced on a MiSeq instrument. Sequencing data were subsequently processed with MOIRAI and custom R scripts as described above.

## Data availability

nanoCAGE sequence data: Zenodo 1680999 (https://doi.org/10.5281/zenodo.1680999)

nanoCAGE sequence alignments: Zenodo 1683162 (https://doi.org/10.5281/zenodo.1683162)

Single-cell sequence data: Zenodo 250156 (https://doi.org/10.5281/zenodo.250156)

Single-cell sequence alignments: Zenodo 3340196 (https://doi.org/10.5281/zenodo.3340196)

## Acknowledgements

We thank Matthias Harbers and Anthony Genot for critical comments on the manuscript and our funders: Grant-In-Aid for Scientific Research (S) 16H06328, RIKEN DGT, RIKEN DGM, RIKEN single-cell project.

## Author contributions (https://casrai.org/credit/)

Conceptualization: CP

Data curation: SP, OA, SK, CP

Formal analysis: SP, OA, SK

Funding acquisition: PC, CP

Investigation: SP, OA, SK, IC

Methodology: CP

Project administration: PC, CP

Resources: IC, HI

Software: SP, CP

Supervision: CP

Validation: SP, OA, SK, CP

Visualization: CP

Writing – original draft: SP, CP

Writing – review & editing: SP, CP

